# Reconstructing the genetic history of Kra-Dai speakers from Thailand

**DOI:** 10.1101/2022.06.30.498332

**Authors:** Piya Changmai, Jan Kočí, Pavel Flegontov

## Abstract

Genetic history of the Thai people and, more generally, speakers of the Kra-Dai languages (also known as Tai-Kadai languages) in Thailand remains a topic of debate. Recently, Kutanan *et al*.^1^ analyzed genome-wide genetic data for dozens of present-day human populations from Thailand and surrounding countries and concluded that the Central Thai, Southern Thai, and Malay from Southern Thailand are genetically continuous with Austroasiatic speakers such as Mon, and thus the advent of Kra-Dai and Austronesian languages to Central and Southern Thailand was overwhelmingly a result of cultural rather than genetic diffusion. We re-analyzed the genetic data reported by Kutanan *et al*.^1^ using an advanced technique for inferring admixture graph models, using autosomal haplotypes, and other methods. We did not reproduce the results by Kutanan *et al*.^1^, and our analyses revealed a more complex picture of the genetic history of Kra-Dai speakers and other populations of Thailand.

## Introduction

Kra-Dai is a language family uniting about 90 languages spoken mainly in Southern China, Laos, Thailand, Vietnam, and Myanmar^2^. It is believed based on evidence from historical linguistics that this language family has been spreading from Southern China in the last two millennia^3^. Limited genome-wide archaeogenetic data that were possible to generate in the tropical climate showed that speakers of another large language family distributed over the same territory, Austroasiatic, are genetically very close to the first farmers in Mainland Southeast Asia (MSEA)^4,5^, but Kra-Dai speakers are not fully genetically continuous with these farmers and likely represent a later wave of migrants to MSEA^5^. Similar to discussion of other large cultural areas, with respect to the area inhabited by Kra-Dai speakers there is a debate around the contribution of cultural vs. demic diffusion to the origin of this area^1,6-8^. Kutanan *et al*.^1^ generated genome-wide genotyping data (on the HumanOrigins SNP array^9^) for 17 present-day Kra-Dai-speaking groups from Thailand and Laos, and for other groups from Thailand. One of the main conclusions of that study is the existence of nearly perfect genetic continuity between Austroasiatic speakers (represented, for example, by Mon) and Central Thai, Southern Thai, and Malay from Southern Thailand. Another scenario for the arrival of the latter three populations to Central and Southern Thailand, that is genetic admixture between the incoming Kra-Dai and Austronesian migrants and the indigenous Austroasiatic population, was not discounted by Kutanan *et al*.^1^, but cultural diffusion was clearly favored as an interpretation for the results of their genetic analyses. These results rely on two lines of evidence: 1) most importantly, admixture graphs fitted to allele frequency data in the form of *f*_*3*_-statistics^9^ and co-modelling Central Thai, Southern Thai, Malay from Southern Thailand, and Mon, in addition to several reference populations, 2) a principal component analysis (PCA) of genetic data^10^, and 3) “ancestry painting” performed with *GlobeTrotter*, a tool relying on autosomal haplotypes^11^.

In this study we re-analyzed the HumanOrigins genotyping data published by Kutanan *et al*.^1^ along with other compatible data for present-day groups from MSEA, East Asia, and South Asia^12-16^. We relied on analytical protocols different from those employed by Kutanan *et al*.^1^ but tailored to the same types of data: 1) *findGraphs*^17^, a tool for exploration of large admixture graph topology spaces and for comparison of admixture graph model fits in a rigorous way (another automated admixture graph inference tool, *AdmixtureBayes*, was used in the original study; see https://github.com/svendvn/AdmixtureBayes); 2) *SOURCEFIND v*.*2*^18^, a haplotype-based tool building an admixture model for a target group from a panel of source proxies (*GlobeTrotter* was used instead in the original study). Our re-analysis does not support the genetic continuity of Austroasiatic speakers with the Thai and Malay groups in Thailand. In addition to a methodological discussion, below we present a fine-grained model for recent ancestry of the Thai and of Kra-Dai speakers in Thailand in general.

## Results and Discussion

### Admixture graph models of genetic history

Kutanan *et al*.^1^ presented two admixture graph models relevant for the central question of genetic continuity between Austroasiatic and Kra-Dai speakers in Thailand. The simpler model (in Fig. 6C of that study) fitting the data well (with the worst-fitting *f*-statistic 1.6 standard errors away from the observed value) included Southern Thai and Malay as groups cladal with Mon, and Central Thai got 22% of their ancestry from another East Asian source according to that model. The more complex model (in Suppl. Fig. 19 of that study) fitted the data poorly (it had the worst *f*-statistic residual of 4.1 SE) and was interpreted as largely supporting the simpler model. According to the complex model, Southern Thai, Central Thai, and Malay from Southern Thailand are essentially cladal with two Austroasiatic-speaking groups (Mon and Cambodians) but differ from them slightly in the proportion of South Asian ancestry or Atayal-related ancestry in the case of Malay. In other words, the sources of East Asian ancestry in Mon, Cambodians, and Thai are the same (see the complex published graph in Fig. 1d). These two admixture graph topologies were inferred automatically using the *AdmixtureBayes* tool (https://github.com/svendvn/AdmixtureBayes), and no alternative models were shown or discussed. Notably, the topological details of these two graphs were used by Kutanan *et al*.^1^ as primary evidence to support a major conclusion: the genetic continuity between Austroasiatic groups and Thai or Malay^1^.

**Fig. 1.**
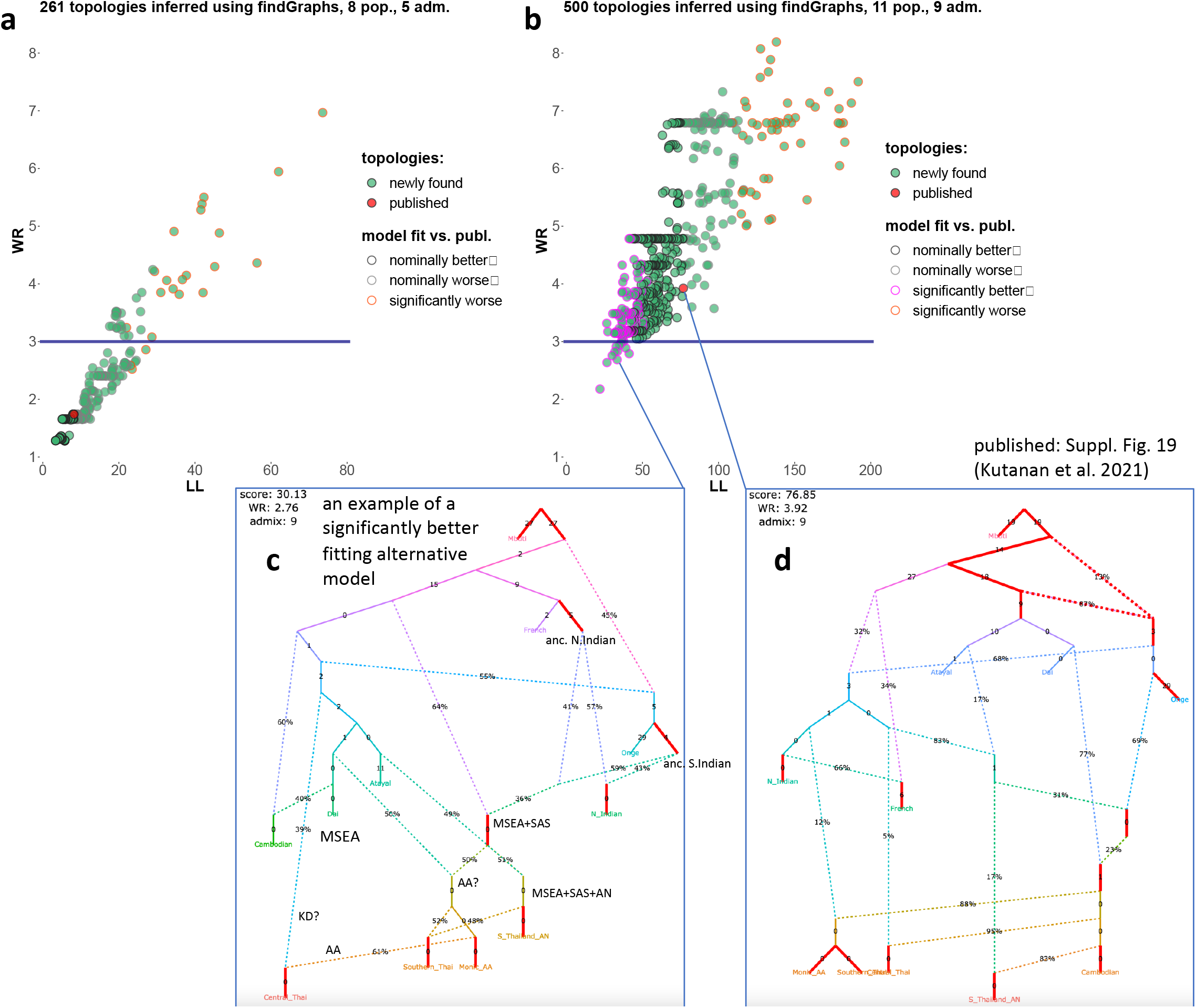
Fits of the newly found and published admixture graph topologies to the HumanOrigins autosomal genetic data. Each distinct topology is visualized as a dot in the space of two model fit metrics: log-likelihood score (LL) and worst *f*-statistic residual (WR), with the published model highlighted in red. Results for the simpler complexity class (8 groups and 5 admixture events) are shown in panel **a**, and results for the complex graphs (11 groups and 9 admixture events) are shown in panel **b**. Results of model fit comparison tests on bootstrap replicates of the dataset^17^ are represented by different border colors according to the legend. For example, models fitting significantly better than the published model are represented by circles with magenta borders in panel **b**. The fitted complex published model and its fit metrics (log-likelihood score and WR) are visualized in panel **d**, and an alternative complex model chosen as an example and its fit metrics are shown in panel **c**. Some edges of the alternative model can be interpreted as ancient populations attested or inferred in the archaeogenetic literature, and those are labeled in panel **c**. Model parameters (admixture proportions and edge lengths measured in units of genetic drift) that cannot be estimated independently are highlighted in red. An algorithm for finding such parameters was introduced by Maier *et al*.^17^ The Malay group from Thailand is labelled as “S_Thailand_AN” on the graphs.

We argue that using admixture graph models in this way, i.e., to support very specific statements about demographic history, is an exercise that is fraught with problems on many levels. These conceptual problems are discussed in detail by Maier *et al*.^17^, where novel approaches for admixture graph inference are introduced. Here we revisited the same population sets and graph complexities (the number of admixture events allowed) that were used by Kutanan *et al*.^1^ and explored these spaces of graph topologies with an automated tool, *findGraphs*^17^, that finds models corresponding to local fit optima in these spaces. In other words, we attempted to find alternative well-fitting models for the simple and complex graphs presented by Kutanan *et al*.^1^, based on a very similar set of SNPs (HumanOrigins) and having the same population composition, the same complexity, and the same outgroup populations as in the original study. The *findGraphs* algorithm is seeded by a random graph of a given complexity and satisfying given constraints, and applies various graph-modification procedures iteratively, attempting to find a local optimum in the graph fit space (see Maier *et al*.^17^ for details). For each graph complexity class, the algorithm was started 500 times from random graphs, and for simplicity, only one inferred graph (best-fitting according to the log-likelihood score) was taken from each such run (see Methods for details). Fits of the resulting sets of distinct alternative topologies to the genetic data are visualized in Fig. 1.

The first problem of the admixture graph framework becomes obvious when inspecting Fig. 1a,b: even a shallow exploration of the enormous spaces of alternative topologies reveals that dozens to hundreds of topologies fit the data approximately equally well. Two metrics are used in the literature for estimating fits of admixture graphs to *f*-statistic data, and those are worst *f*-statistic residuals (WR), also referred to as Z-scores and measured in standard errors (SE), or log-likelihood scores (LL) that take into account all *f*-statistics. Here we placed the newly inferred and the published models in the space of both metrics that are relatively well correlated (Fig. 1a,b).

Comparing fits of alternative admixture graph models in a statistically rigorous way is needed for any large-scale model exploration, and an algorithm for this purpose was introduced by Maier *et al*.^17^ The essence of this algorithm is fitting two alternative models on a set of bootstrap-resampled replicates of genetic data (this is easy to implement since for calculating SEs of *f*-statistics SNPs are divided into blocks based on genetic or physical distance) and comparing the resulting distributions of LL scores (see Maier *et al*.^17^ for details). Unlike previous methods for comparing fits of admixture graph models^19,20^, this method takes stochasticity in evolution of unlinked SNPs into account and has no assumptions about the number of independent model parameters (which in the case of admixture graphs is not trivial to estimate)^17^. Relying on this bootstrap-based model comparison approach, we found that hundreds of alternative models (matching both the simple and complex graphs in complexity) have fits to the data that are not significantly different from that of the published model (Fig. 1a,b). We note that the same constraints on the graph topology were applied as in the original study: French or Mbuti were assigned as an outgroup. Even a shallow exploration of both graph spaces found hundreds of models that fit the data as good as the published ones, and deeper exploration (performing more *findGraphs* runs and/or extracting more graphs from each run) is guaranteed to deliver further and further models of this kind^17^.

A question arises: is it justified to put a lot of weight on a particular graph and derive historical interpretations of its topology if hundreds of diverse topologies fit the data equally well? This is a key point discussed by Maier *et al*.^17^ when revisiting admixture graphs from eight published studies. For instance, it was found that there are at least several models fitting the data significantly better than the admixture graph for East Asians used to support a key conclusion by Wang *et al*.^13^, and the alternative models do not support the conclusion. These alternative models were found even after applying multiple topological constraints (guided by archaeology, linguistics, and other genetic studies) that Wang *et al*.^13^ relied on when constructing their graph manually^17^.

What are the reasons for this high topological diversity among well-fitting models that was shown to be ubiquitous for admixture graph spaces explored in the literature?^17^ First, there are inherent limitations of *f*-statistics related to directionality of gene flow: distinct graph topologies are known to yield identical *f*-statistics (see, for instance, Prüfer *et al*.^21^). Second, overfitting becomes a problem if too many admixture events are allowed. Overfitting was shown to be a common problem of admixture graphs reported in the literature^17^. Third, diversity of well-fitting topologies may result from a lack of reference populations that are differentially related to populations of interest and are needed for constraining the models^17^.

We believe that the latter point is especially relevant for interpreting the admixture graph results by Kutanan *et al*.^1^ We note that all the MSEA groups included in the published graphs (Cambodians, Mon, Central and Southern Thai, Malay) are separated by very short genetic drift edges: the lengths of these edges are very close to 0 in the case of the simple (see Fig. 6C in the original study) and complex (Fig. 1d) published graphs and all the alternative models we explored (see an example in Fig. 1c). This suggests that reference groups that could be instrumental in distinguishing the populations of interest (due to their differential relatedness to them) are lacking in the models. The simple eight-population graph is especially problematic in this respect since the only East Asian group included that lives outside of the region of interest (MSEA) is Atayal from Taiwan, and no Tibeto-Burman-speaking or Kra-Dai-speaking proxies for potential ancestry sources were included. For instance, a Tibeto-Burman-related ancestry component was detected in Austroasiatic-speaking Mon by Kutanan *et al*.^1^ using methods other than admixture graphs, and also by Changmai *et al*.^14^ Given these results, the lack of a Tibeto-Burman reference population (and of other key reference groups) in both the simple and complex graphs from Kutanan *et al*.^1^ probably makes these admixture graph systems unconstrained. Therefore, it is not surprising that hundreds of simple topologies fit the data well in absolute terms (WR < 3 SE) and fit the data as good as the published model (Fig. 1a).

Since the complex graphs are more constrained than the simple ones (likely due to the inclusion of Kra-Dai-speaking Dai from Southern China), the alternative models we found are more differentiated according to their fits to the data (Fig. 1b). The published model on our dataset has a fit (WR = 3.9 SE) that is very close to that reported in the original study (WR = 4.1 SE), and such a fit is considered poor by convention since it exceeds 3 SE. We found 11 complex topologies (with 11 groups and 9 admixture events) that fit the data well in absolute terms (WR < 3 SE) and, moreover, that fit the data significantly better than the published topology (with two-tailed empirical model-comparison *p*-values < 0.05). One such topology is shown as an example in Fig. 1c. As discussed in Maier *et al*.^17^, inference of demographic history in the admixture graph framework has important limitations even if best-practice protocols introduced in that study are adhered to. For instance, it is unknown what parsimony level (the number of admixture events) is optimal for inferring true history, and changing graph complexity can dramatically change the pattern of topologies that fit the data, and hence their historical interpretation^17^. As discussed above, the outcomes of a model inference protocol also depend a lot on the choice of groups included in the model^17^. For these reasons we did not believe that any well-fitting model found by us is accurate; on the contrary, we believe that all of them are wrong in one way or another. However, the model shown in Fig. 1c has several features that match archaeogenetic results reported in the literature and derived using various methods other than admixture graphs: 1) Indians are derived from Ancient North Indians of West Eurasian origin and Ancient South Indians related to the Andamanese^22,23^; 2) there is a fraction of Atayal (Austronesian)-related ancestry in Malay from Southern Thailand, who are Austronesian speakers^14^; 3) there is South Asian (Indian) ancestry in nearly all MSEA groups included in the model^1,14^.

Importantly, the alternative model shown in Fig. 1c (and other alternative models we found) contradict the key conclusion by Kutanan *et al*.^1^, that is the nearly perfect genetic continuity between the Austroasiatic (Mon) and Thai and Malay groups in Thailand. If we interpret the graph node marked as “MSEA+SAS” (Fig. 1c) as an Austroasiatic-speaking group with Indian admixture similar in its ancestry composition to Mon^1,14^, then the Malay group from Thailand gets 49% of ancestry from an Austronesian Atayal-related source, Southern Thai get 48% of their ancestry from Malay, and Central Thai get 39% of their ancestry from an unidentified East Asian source (Fig. 1c). This topology and inferred admixture proportions are hardly compatible with the scenario of genetic continuity between indigenous Austroasiatic speakers represented by Mon and Thai or Malay people. We stress that although the alternative model presented in this study fits the data well and significantly better than the published one, we do not claim it to be fully accurate. We believe that for disproving a claim relying on a particular admixture graph topology, even a single well-fitting historically plausible topology serving as a counterexample is enough.

### Inference of recent ancestry based on autosomal haplotypes and other evidence

Another result by Kutanan *et al*.^1^ supporting the genetic continuity hypothesis is inference of recent ancestry with *GlobeTrotter*^11^. According to this analysis, the Mon, Central Thai, Southern Thai, and Malay from Thailand were inferred to have similar ancestry profiles (proportions of ancestry derived from a panel of potential sources according to the model), and at least 50% of ancestry in these four groups was contributed by a source most closely related to Austroasiatic-speaking Kinh (Vietnamese, see Fig. 6A in the original paper). These results (the similarity of “ancestry painting” profiles for Mon and Thai or Malay and the high proportion of Kinh-related ancestry in all these groups) were interpreted as supporting the genetic continuity hypothesis for Austroasiatic speakers and Thai and Malay.

*SOURCEFIND*^18^ was introduced by the team that developed *ChromoPainter*^11,24^ and *GlobeTrotter*^11^, and unlike the latter software, which is mainly aimed at inference of admixture dates, *SOURCEFIND* is aimed specifically at inferring complex mixture models (proportions of admixture). This tool implements a mixture model distinct from that used in GlobeTrotter, and this model demonstrates better performance on simulated data^18^. For this reason, we decided to reanalyze the data by Kutanan *et al*.^1^ with *SOURCEFIND*. Our analysis was focused on 15 Kra-Dai-speaking groups from Thailand (Fig. 2, Suppl. Table 2). Three Austroasiatic-speaking (Bru, Khmu, and Palaung), one Hmong-Mien-speaking (Hmong Daw), and one Sino-Tibetan-speaking group (Karen Padaung) were chosen as controls. Most other MSEA groups and selected East Asian and South Asian groups from our dataset (Suppl. Table 1) were used as potential ancestry source proxies. Unlike *GlobeTrotter, SOURCEFIND* identifies source proxies whose contribution is distinguishable from noise and uses only those in constructing a mixture model. We show ancestry proportions for sources accounting for at least 1% of the genome in any target group in Suppl. Table 2 and for major sources (>10% of the genome in at least one group) in Fig. 2. We also note that another difference in the approaches used here and by Kutanan *et al*.^1^ is the composition of the panel of potential source proxies (Suppl. Table 1 and Fig. 6A in Kutanan *et al*.^1^). For instance, no Kra-Dai speakers from Laos and Southern China (except for Dai) were included in this panel by Kutanan *et al*.^1^

**Fig. 2.**
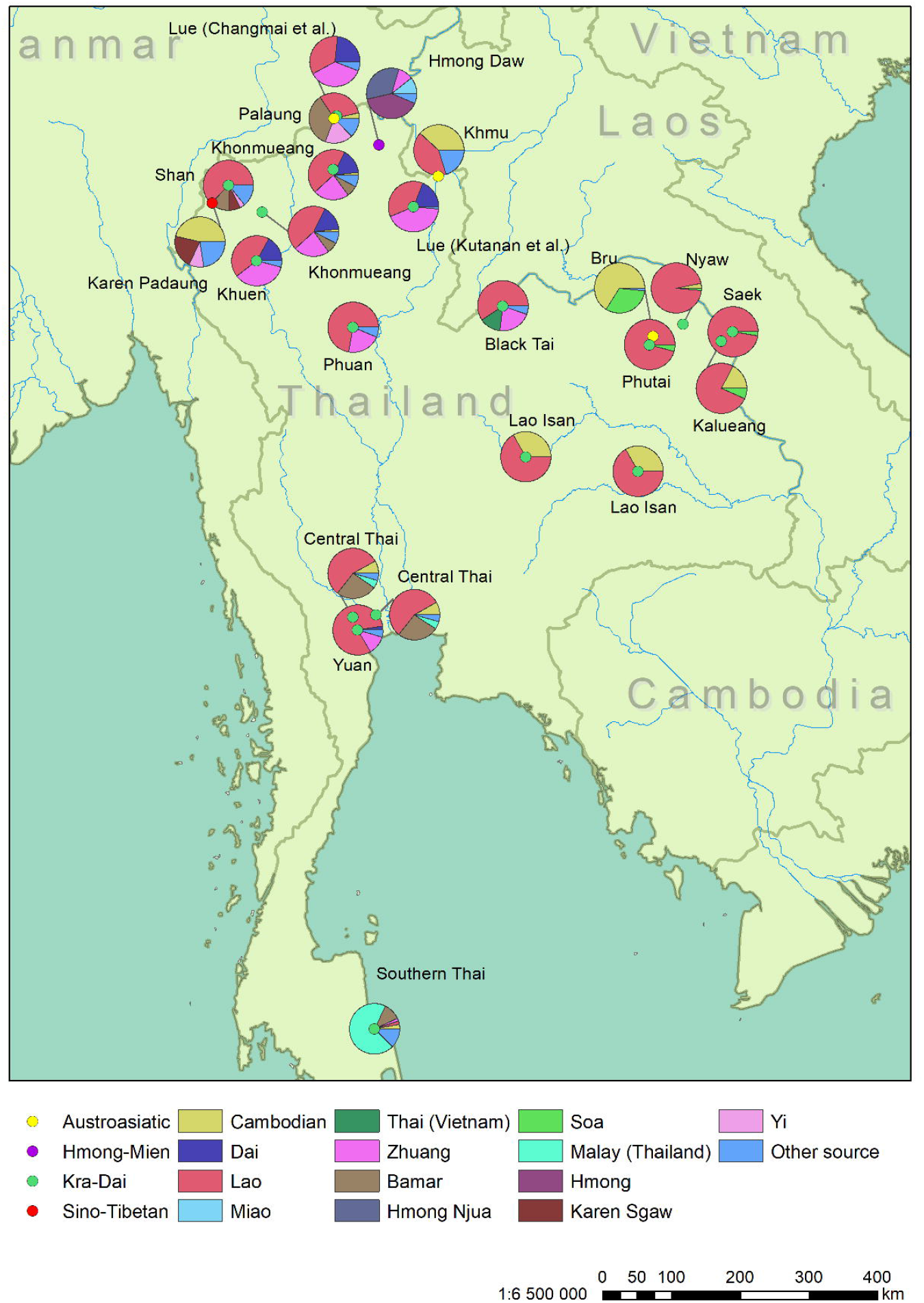
Sources of recent ancestry as inferred with *SOURCEFIND v*.*2* in groups from Thailand. Here only sources contributing >10% to at least one target group are visualized (for full results see Suppl. Table 2). Locations of the groups on the map are shown with circles, and those are colored according to language affiliation. Ancestry composition is illustrated using pie charts.

According to our mixture model (Fig. 2, Suppl. Table 2), a predominant ancestry component in most Kra-Dai speakers from Thailand is not Kinh-related, but Lao-related (a Lao group from Laos was used as a source proxy), although Kinh was also included in the panel of potential source proxies (Suppl. Table 1). The fraction of Lao-related ancestry reached 95% in Kra-Dai speakers from the Northeast of Thailand near the Laos border. In contrast, the fraction of Kinh-related ancestry reached at most 4.4% in any of the target groups (Suppl. Table 2). Genetic contribution from a Mon-related Austroasiatic source was also negligible in Kra-Dai speakers according to our model, 1% at most (Suppl. Table 2). Bru, another group from Northeastern Thailand, demonstrates a strikingly different pattern, with <1% of Lao-related ancestry (Fig. 2, Suppl. Table 2). Bru is a relatively isolated Austroasiatic-speaking group, and thus the patterns of recent ancestry inferred with *SOURCEFIND* are influenced not only by geography. The Lao-related ancestry component accounts for >50% of ancestry in the Central Thai, but is negligible in the Southern Thai. Although an Austroasiatic genetic component (Cambodian-related) contributes to some Kra-Dai groups in Thailand according to our model (Fig. 2, for example, 7.5% in the Central Thai), in none of these groups it accounts for more than 32% of ancestry. In addition to Laos, we were able to trace genetic connections to Kra-Dai-speaking groups in Southern China: the Zhuang-related component accounted for up to 41% of ancestry in Kra-Dai speakers, but only in the North of Thailand (Fig. 2). Remarkably, it was also detected at 11.5% in the Yuan group from Central Thailand who were resettled from Northern Thailand about 200 years ago^25^.

In agreement with our admixture graph model for Southern Thai (Fig. 1c), where 48% of their ancestry is derived from a Malay (Austronesian)-related source, this groups was modelled by *SOURCEFIND* as having 66% of their ancestry derived from the same source. We stress that the Malay group involved^1^ was sampled in Thailand, thus this result probably reflects bidirectional gene flow between Malay-speaking and Thai-speaking groups in Southern Thailand^26^.

A surprising result of our inference of recent ancestry with *SOURCEFIND* is a large proportion of Bamar-related ancestry in the Central (24%) and Southern Thai (11%) (Fig. 2). This ancestry component was also detected in two groups (Palaung and Shan) located close to the border with Myanmar (Fig. 2), and in that case the result correlates with geography. Our *SOURCEFIND* analysis (Suppl. Table 2) did not detect appreciable South Asian ancestry in the target groups, but we believe that the signal was obscured by the presence of groups with substantial South Asian admixture among the source proxies (Bamar and Cambodians)^14^.

Thus, to extend our previous mapping of South Asian ancestry in present-day MSEA^14^, we performed dedicated analyses with *GlobeTrotter* (Suppl. Table 3), *ALDER* (Suppl. Table 4), and “admixture” *f*_*3*_-statistics (Suppl. Table 5). We applied *GlobeTrotter* to exactly the same dataset as was analyzed with *SOURCEFIND*, and substantial South Asian admixture was detected in the following groups: in Kra-Dai-speaking Southern Thai (31%), Central Thai (24%), Lao Isan (22%), and Khonmueang (16%), and in Austroasiatic-speaking Palaung (25%) (Suppl. Table 3). South Asian ancestry in all five groups was supported not only by *GlobeTrotter*, but also by *ALDER* based on linkage disequilibrium decay (Suppl. Table 4), and by “admixture” *f*_*3*_-statistics^9^ of the type *f*_*3*_(MSEA; SEA, South Asian) based on allele frequency correlations (Suppl. Table 5). In the case of *ALDER* and *f*_*3*_-statistics, all possible pairs of source proxies “East or Southeast Asian + South Asian” were tested.

## Conclusions

We aimed at testing a historical scenario recently proposed by Kutanan *et al*.^1^, namely that the Central Thai, Southern Thai, and Malay from Thailand are genetically continuous with indigenous Austroasiatic speakers such as Mon. We demonstrated that this simple model favored by Kutanan *et al*.^1^ fits allele frequency correlation data (*f*-statistics) significantly worse that some alternative admixture graph models we found relying on a new tool for exploring admixture graph fit spaces^17^. We also demonstrated that the other line of evidence invoked by Kutanan *et al*.^1^, namely complex mixture models based on autosomal haplotypes, depends critically on details of the modelling algorithm (*GlobeTrotter* vs. *SOURCEFIND*) and on the composition of the panel of source proxies. Our mixture models inferred with *SOURCEFIND* suggest a rich history of Kra-Dai speakers in Thailand, with several ancestry components correlated with geography or linguistic affiliation. In general, our models, both admixture graphs and *SOURCEFIND*, support substantial Austroasiatic-related admixture in Kra-Dai speakers from Thailand, but they reject the model of genetic continuity between these major populations.

Kutanan *et al*.^1^ detected substantial South Asian admixture in the Central Thai, Southern Thai, and Malay from Thailand, but not in other newly reported groups. We extended this result to few other groups from Thailand reported by Kutanan *et al*.^1^: Kra-Dai-speaking Lao Isan and Khonmueang, and Austroasiatic-speaking Palaung. Combining these results with our previous study focused on South Asian admixture across MSEA^14^, we conclude that this ancestry is common in the region, but far from universal. South Asian ancestry is likely restricted to populations that were involved in the formation of early states in MSEA influenced by Indian culture such as Funan in Cambodia and Dvaravati in Thailand^14^.

## Methods

### Assembling the dataset

We used exclusively published diploid genotyping data generated on the Affymetrix HumanOrigins SNP array^9^ mainly in the following studies: Kutanan *et al*.^1^, Liu *et al*.^12^, Wang *et al*.^13^, Changmai *et al*.^14^, Lazaridis *et al*.^15^, Nakatsuka *et al*.^16^ For a list of individuals, groups, their linguistic affiliations, and data sources see Suppl. Table 1. All our work relied on a set of 574,131 autosomal HumanOrigins SNPs identical to that used in Changmai *et al*.^14^

### Exploring admixture graph topology spaces

All our work with *f*-statistics and methods relying on them was done using the *ADMIXTOOLS 2* package^17^ (https://uqrmaie1.github.io/admixtools/). To calculate *f*_*3*_-statistics needed for fitting admixture graph models, we first used the “*extract_f2*” function with the “*maxmiss*” argument set at 0, which corresponds to the “*useallsnps: NO*” setting in the classic *ADMIXTOOLS*^9^. It means that no missing data are allowed (at the level of populations) in the specified set of populations for which pairwise *f*_*2*_-statistics are calculated. The “*blgsize*” argument sets the SNP block size in Morgans, and we used the default value of 0.05 (5 cM). Since all groups involved in the admixture graph modelling included more than one individual, and diploid variant calls were available for all individuals, the “*adjust_pseudohaploid*” and “*minac2*” arguments were set to “*FALSE*”^17^. The “*extract_f2*” function calculates *f*_*2*_-statistics for all pairs of groups per each SNP block, and those are used by the “*find_graphs*” and “*qpgraph*” functions for calculating *f*_*3*_-statistics as linear sums of *f*_*2*_-statistics^9^. In the absence of missing data (and no missing data weas allowed at the level of groups) the linear sums should be unbiased^17^.

We used the “*find_graphs*” function from the *ADMIXTOOLS 2* package for finding multiple alternative well-fitting topologies. We worked on the sets of groups and individuals that were identical to those used by Kutanan *et al*.^1^ for constructing their admixture graphs presented in Fig. 6C (8 groups and 5 admixture events) and Suppl. Fig. 19 (11 groups and 9 admixture events). We did not modify the graph complexity (the number of admixture events) either and kept the outgroups used by Kutanan *et al*.^1^: French for the simpler graph and Mbuti for the complex graph. The characteristics of the datasets used for admixture graph fitting in our study are as follows: 1) graphs of the “simple” complexity class were based on 8 groups, 177 individuals (Suppl. Table 1), and 456,719 sites polymorphic in this set of groups and having no missing data at the group level; 2) graphs of the “complex” class were based on 11 groups, 207 individuals (Suppl. Table 1), and 501,703 sites polymorphic in this set of groups and having no missing data at the group level.

For each graph complexity class, the *findGraphs* algorithm was started 500 times independently, seeded by random graphs with a specified number of admixture events (5 or 9) and a specified outgroup (French or Mbuti). Random graphs were generated with the “*random_admixturegraph*” function. The settings of the *findGraphs* algorithm were identical to those presented in Maier *et al*.^17^ (see the Methods section in that preprint), and French or Mbuti were specified as outgroups at this topology optimization step too. From each *findGraphs* run, one best-fitting topology (i.e., the highest-ranking topology according to the log-likelihood score) was extracted, and a set of non-redundant topologies was constructed from all the runs. Fits of these topologies to the data (log-likelihood scores and the worst *f*-statistic residuals) were plotted, and best-fitting topologies were inspected manually for features that are important for historical interpretations. The published admixture graph topologies (Fig. 6C and Suppl. Fig. 19 from Kutanan *et al*.^1^) were fitted to the same per-block *f*_*2*_-statistic data using the “*qpgraph*” function with the following settings: “*numstart=100, diag = 0*.*0001, return_fstats=TRUE*”.

To find out if newly found admixture graph models fit the data significantly better or worse than the published ones, we used a bootstrap-based model comparison algorithm developed by Maier *et al*.^17^ Five hundred bootstrap replicates of the two SNP block datasets, corresponding to the simple and complex published graphs, were generated (with the 5 cM block size). The algorithm reports empirical two-tailed *p*-values, 0.05 was used as a *p*-value threshold, and the settings of the algorithm were identical to those used by Maier *et al*.^17^

### Methods based on autosomal haplotypes

We phased a world-wide dataset of 3,945 individuals (compiled from published sources) using *SHAPEIT*

*v*.*2 (r900)*^27^ with 1000 Genomes Phase 3 genetic maps^28^. We then ran *ChromoPainter v*.*2*^11,24^ to generate inputs for *SOURCEFIND v*.*2*^18^ and *fastGLOBETROTTER*^29^. We selected 75 surrogates and 20 target populations (14 Kra-Dai-speaking groups from Thailand for whom the data were reported by Kutanan *et al*.^1^, one Kra-Dai-speaking group from Thailand for whom the data were reported by Changmai *et al*.^14^, and 5 control groups from Thailand speaking other languages) (see a list of populations involved in Suppl. Table 1). We ran *ChromoPainter v*.*2* assigning all the surrogates as donors and recipients, but the target populations were assigned as recipients only. This means that target populations receive haplotypes only from surrogates, but not from their own population nor other target populations. The other details of the *ChromoPainter v*.*2* protocol exactly followed those presented by Changmai *et al*.^14^

The settings of the *SOURCEFIND* algorithm used for inferring complex mixture models for the same set of 20 target groups were identical to those used by Changmai *et al*.^14^, and all the 75 surrogates were used as a panel of potential sources from which the algorithm constructed mixture models for each target. The settings of the *fastGLOBETROTTER* algorithm were also identical to those used by Changmai *et al*.^14^, and all the 75 surrogates were used for inferring best-fitting admixture models and estimating admixture dates.

### Fitting admixture models to linkage disequilibrium decay curves

We used the *ALDER* tool^30^ with the default settings for fitting two-way admixture models of the type “East or Southeast Asian group + South Asian group” for the set of 20 target groups. For a list of groups involved see Suppl. Table 4.

### f_3_-statistics

Statistics of the type *f*_*3*_(one of the 20 target groups; an East or Southeast Asian group, a South Asian group) were calculated using the “*qp3pop*” function of the *ADMIXTOOLS 2* package. For each triplet of groups, no missing data was allowed at the group level (the default setting). *f*_*3*_-statistics were calculated directly from the genotype data, without *f*_*2*_-statistics as an intermediate. For a list of groups involved see Suppl. Table 5.

## Acknowledgments

This work was supported by the Czech Ministry of Education, Youth and Sports: 1) Inter-Excellence program, project #LTAUSA18153; 2) program ERC CZ, project no. LL2103; 3) Large Infrastructures for Research, Experimental Development and Innovations project “IT4Innovations National Supercomputing Center – LM2015070”. P.F. was also supported by a subsidy from the Russian federal budget (project No. 075-15-2019-1879 “From paleogenetics to cultural anthropology: a comprehensive interdisciplinary study of the traditions of the peoples of transboundary regions: migration, intercultural interaction and worldview”).

## Author contributions

P.C. designed and P.F. supervised the study. P.C., J.K., and P.F. analyzed the data. P.C. and P.F. drafted the manuscript with additional input from J.K.

## Supplemental Tables

**Suppl. Table 1**. Composition of the dataset: groups and their sizes, linguistic affiliations, studies where the data were first reported, and their involvement in analyses in this study.

**Suppl. Table 2**. Complex admixture models for 20 target groups inferred with *SOURCEFIND* aimed at detecting recent ancestry. Data for nineteen groups were published by Kutanan *et al*.^1^, and data for a Lue group were published by Changmai *et al*.^14^ Only source proxies contributing at least 1% of ancestry in any target group are shown in this table.

**Suppl. Table 3**. Most likely two-way or multi-way admixture models inferred with *GlobeTrotter* (and their fits to the data and inferred admixture dates) for the same target groups and the same proxy sources that were analyzed with *SOURCEFIND*.

**Suppl. Table 4**. Inference of two-way admixture models and admixture dates with *ALDER*. The set of 20 target groups was analyzed (the same as in Suppl. Tables 2, 3, and 5), but in contrast to the *GlobeTrotter* approach, only pairs of source proxies composed of an East or Southeast Asian group and a South Asian group were tested. Only successfully fitted models are shown in the table.

**Suppl. Table 5**. “Admixture” *f*_*3*_*-*statistics of the type *f*_*3*_(MSEA; SEA, South Asian). The set of 20 target groups was analyzed (the same as in Suppl. Tables 2, 3, and 4). A significantly negative value (Z-score < 3 SE) is a proof that the target group is admixed between sources related closely or distantly to the two other groups^9^.

